# Force-Dependent Cell-Cell Adhesion Dynamics in a Stochastic Regime for Cancer Invasion

**DOI:** 10.64898/2026.03.11.710757

**Authors:** Sascha Schultz, Dimitrios Katsaounis, Nikolaos Sfakianakis

**Affiliations:** School of Mathematics and Statistics, University of St Andrews, North Haugh, St Andrews, KY16 9SS, Scotland, UK; Institut für Geometrie und Praktische Mathematik, RWTH Aachen University, Im Süsterfeld 2, Aachen, 52072, Germany

## Abstract

Cell-cell adhesion is a key regulator of cancer invasion. In this work, we extend a pre-existing individual based cancer invasion model by introducing a stochastic representation of N-cadherin-mediated adhesion, where the lifetime of a cell-cell bond depends on the pulling force acting on the bond. Using experimental data, we derive expressions for the mean and standard deviation of N-cadherin bond lifetimes and fit them to Gamma distributions, enabling their treatment as force-dependent random variables. These distributions are then used to modify the diffusion coefficient of mesenchymal cancer cells. The model predicts reduced random motility with increasing adhesion and incorporates a dynamic transition between catch- and slip-bond behaviour. Along with this model for cell motility, we propose a preliminary physical framework, that can be used to model pattern formation as a result of the new adhesion mechanic.

## 1. Introduction

Cancer remains a major global health burden, contributing to approximately 15% of worldwide mortality [1, 2]. While benign tumours are typically confined and rarely life-threatening, malignant tumours acquire genetic and epigenetic phenotypes that enable them to invade adjacent tissues and disseminate to distant sites, posing a serious threat to patient survival [3, 4]. Throughout this work, the focus will be placed on such malignant tumours, primarily characterised by their invasive potential. Cancer invasion, and the subsequent metastasis, is one of the distinct biological hallmarks separating cancer cells from their healthy counterparts. The hallmarks of cancer were introduced by Hanahan & Weinberg [3] and later in [5], underpinning all the necessary characteristics for successful tumour development and progression. In brief, cancer may be understood as a heterogeneous population of cells with the ability to sustain proliferation, evade inhibitory external cues, suppress apoptotic pathways, and induce angiogenesis to ensure adequate supply of oxygen and nutrients. Crucially, these cells invade adjacent tissue structures and, in many cases, disseminate to distant anatomical sites. In this work, we focus on the invasive behaviour of malignant tumours, with an emphasis on how cell-cell adhesion influences the patterns and dynamics of tissue infiltration.

To place the phenomenon of tissue invasion in context, we outline the broader sequence of events underlying cancer progression. We focus, in particular, on carcinomas which are tumours arising from epithelial cells that have acquired a series of mutations that confer malignant potential [6]. These *epithelial-like cancer cells* (ECCs) initially over-proliferate, giving rise to a primary tumour. At this early stage, the tumour consists primarily of ECCs. These cells collectively secrete pro-angiogenic factors that stimulate the formation of new blood vessels, ensuring a sustained supply of oxygen and nutrients and thereby supporting the continuous growth of the tumour [7]. As the tumour evolves, ECCs may undergo further phenotypic changes via the *epithelial-mesenchymal transition* (EMT) [8], resulting in the emergence of *mesenchymal-like cancer cells* (MCCs). These two populations, ECCs and MCCs, differ significantly in their functional properties. Although ECCs retain a high proliferative capacity, their strong cell-cell and weak cell-matrix adhesion forces limit their motility, [9]. On the contrary, MCCs downregulate intercellular adhesion molecules during EMT, rendering them less proliferative but highly motile. This altered adhesive profile enables MCCs to detach from the primary tumour mass and invade the surrounding tissue, thereby initiating the invasion-metastasis cascade [10, 11, 12].

The aim of the present study is to investigate and incorporate the influence of cell-cell adhesion on MCC migration in a mathematical framework for cancer invasion. Firstly, we draw on experimental data quantifying the lifetimes of adhesion bonds under varying levels of mechanical load. These data will inform the construction of fitted probability distributions, allowing us to model the lifetimes of key adhesion molecules as random variables with known distributional properties. Then, the fitted results will be used to update an individual based model of MCC invasion, proposed in [13], that is presented in Section 2.

## 2. The Base Model

The migration of cancer cells has been thoroughly investigated with a variety of mathematical methods where the models used can be separated (although not exclusively) into two main categories: macroscopic models of *Partial Differential Equations* (PDEs) that describe the evolution of the density of the cancer population [14, 15, 16], and individual-based models that describe the invasion of each cell inside the tissue [17, 18, 19, 20, 21, 22]. Macroscopic approaches have been extended to incorporate multiscale and nonlocal effects [23, 24, 25, 26], while individual based models include stochastic particle systems and mean-field limits [27, 28, 29] and detailed computational mechanics [30], as well as recent developments tailored to cancer invasion dynamics [31, 32, 33].

Furthermore, multiscale models have been developed that couple individual-based descriptions for cancer cells with macroscopic density formulations for the remaining biological components involved in the migration process [34, 35, 36, 37]. Related hybrid approaches have also been proposed in [38, 39, 40, 41, 42], where discrete and continuum components are dynamically linked. In addition, hybrid models in which micro- and macroscopic descriptions for cancer cells are coupled for the spatio-temporal growth and migration of the tumour have been employed in [43, 44, 13].

Despite this extensive body of work, the explicit role of cell-cell adhesion in the migration of MCCs remains comparatively less explored within the individual-based stochastic frameworks. Experimental and theoretical studies have emphasised the importance of mechanical coupling, force transmission, and collective coordination during cancer cell migration [45, 46, 47, 21], and recent experimental evidence has further highlighted the role of adhesion-mediated signalling in cooperative metastatic progression [48].

Nevertheless, most existing modelling approaches incorporate cell-cell interactions either through collision-mechanisms or through nonlocal aggregation forces [49, 50, 51, 52, 53, 54], without explicitly linking adhesion dynamics to stochastic motility mechanisms.

Motivated by these observations, we study here the role of adhesion forces among highly migratory MCCs and their impact on stochastic cell motility. In particular, we extend the individual based model, proposed in [43] and further developed in [31, 13], which describes the movement of the cells via *stochastic differential equations* (SDEs) involving drift and a diffusion terms, and which reads as follows:

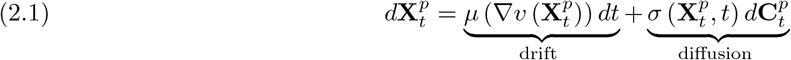

where 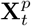 denotes the position of cell *p* at time *t*, and 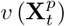 denotes the density of the *extracellular matrix* (ECM) at the position of cell *p* at time *t*. The drift term *µ* depends directly on the density of the ECM, representing that MCCs migrate to regions of higher ECM density. The diffusion coefficient 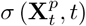 represents the magnitude of the random MCC motility in the ECM, whereas the Compound Poisson Process 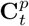 governs the number, timing, and direction of the discrete random ‘jumps’ of the cell. In particular, 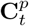 is defined as follows:

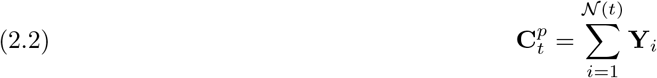

where 𝒩(*t*) is a Poisson process with rate λ *>* 0 and where the 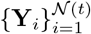 are independent and identically distributed vector-valued random variables. The Poisson process counts the number of independent discrete events that have occur up to time *t*.

The existing model offers a biologically plausible account of cancer invasion, capturing key migratory behaviours. However, it does not incorporate direct physical interactions between individual invading cells. In particular, mechanisms such as cell-cell repulsion and adhesion are currently absent: MCCs may traverse one another without constraint, and their trajectories remain unaffected by neighbouring cells.

In the current work, we will extend the model (2.1)-(2.2) by incorporating intercellular adhesion between MCCs. In particular, we retain the assumption that MCCs primarily migrate towards regions of high ECM density, as in (2.1), and rather than incorporating cell-cell adhesion as direct forces between cells, as it has traditionally been done for individual based models (see e.g. [50, 27, 55]), we assume that adhesion affects both the frequency and magnitude of the random movement of MCCs. Furthermore, the adhesion dynamics will be directly informed by experimental data that measure the lifetime of force-dependent adhesion bonds between cells, which provide estimates for both the rate λ *>* 0 of the Poisson process *𝒩*(*t*) and the magnitude of the 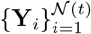 random variables.

The rest of the paper is structured as follows: In Section 3, we present the role of N-cadherins in the creation of adhesive bonds and estimate the mean lifetime of the force-dependent bonds using experimental data. In Section 4, we systematically incorporate the adhesion dynamics into the diffusion coefficient of (2.1) and present the updated individual based model. Finally, in Section 5, we present the results of numerical simulations of the updated model.

## 3. Force-Dependent Bond Lifetimes

In this section we will derive expressions for the lifetime of adhesive bonds based on the pulling force acting on them. For the sake of modelling simplicity, we will assume that these forces are constant over the lifetime of the bond.

Cell-cell adhesion in multicellular systems is mediated by several families of cell-surface proteins, including selectins, cadherins, members of the immunoglobulin superfamily, and integrins [48]. These proteins differ substantially in their mechanical properties, binding specificity, and biological function. In the context of cancer invasion in particular, it is cadherins that mediate stable cell-cell adhesion between MCCs. Cadherins form calcium-dependent bonds capable of sustaining relatively large mechanical loads. Using available experimental measurement data we will model cadherin bond lifetimes and their rupture as inherently stochastic processes.

Of the numerous cadherin subtypes present in human tissues, we restrict attention to the two most widely studied variants: E-cadherin and N-cadherin. E-cadherin forms stronger adhesive bonds and is abundantly expressed in epithelial cells, contributing to the cohesive integrity of ECC populations [57, 58]. During EMT, interactions with fibroblasts lead to downregulation of E-cadherin expression and compensatory upregulation of N-cadherin—the so-called E-to N-cadherin switch—resulting in weakened intercellular adhesion and an increased propensity for MCC motility and metastatic spread [59].

Cadherin-mediated bonds can be classified according to their mechanical response to applied forces. Two principal categories exist: slip bonds, whose lifetimes decrease under force, and catch bonds, whose lifetimes increase under force [60]. Under physiological conditions, catch bonds typically convert into slip bonds once the applied force exceeds a critical threshold, as illustrated in Figure 1.

**Fig. 1:**
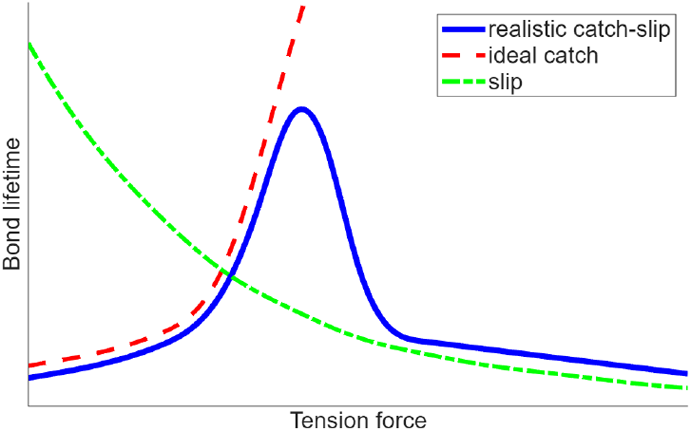
Schematic illustration of bond lifetimes as functions of the applied tension force for different bond types. The figure is inspired by [56].

This force-dependent pattern is closely linked to changes in the dimer conformation of the cadherin molecules: E-cadherin initially forms X-dimers, which function as catch bonds. Over time and with sustained force these X-dimers transition to strand-swap dimers, which behave as slip bonds [61].

While the force-dependent lifetimes of E-cadherin bonds have been verified experimentally, see [60], experimental data on N-cadherin is very scarce, so we shall consider E-cadherin and derive a way of translating this to N-cadherin. Not many experiments have been conducted directly comparing these two bond types, so we will make an educated guess based on two separate studies with similar experimental designs, which considered the number of rupture events at different forces. First, consider the study on N-cadherin shown in Figure 2b, where we see a unimodal right-skewed distribution with its peak at 25pN.

**Fig. 2:**
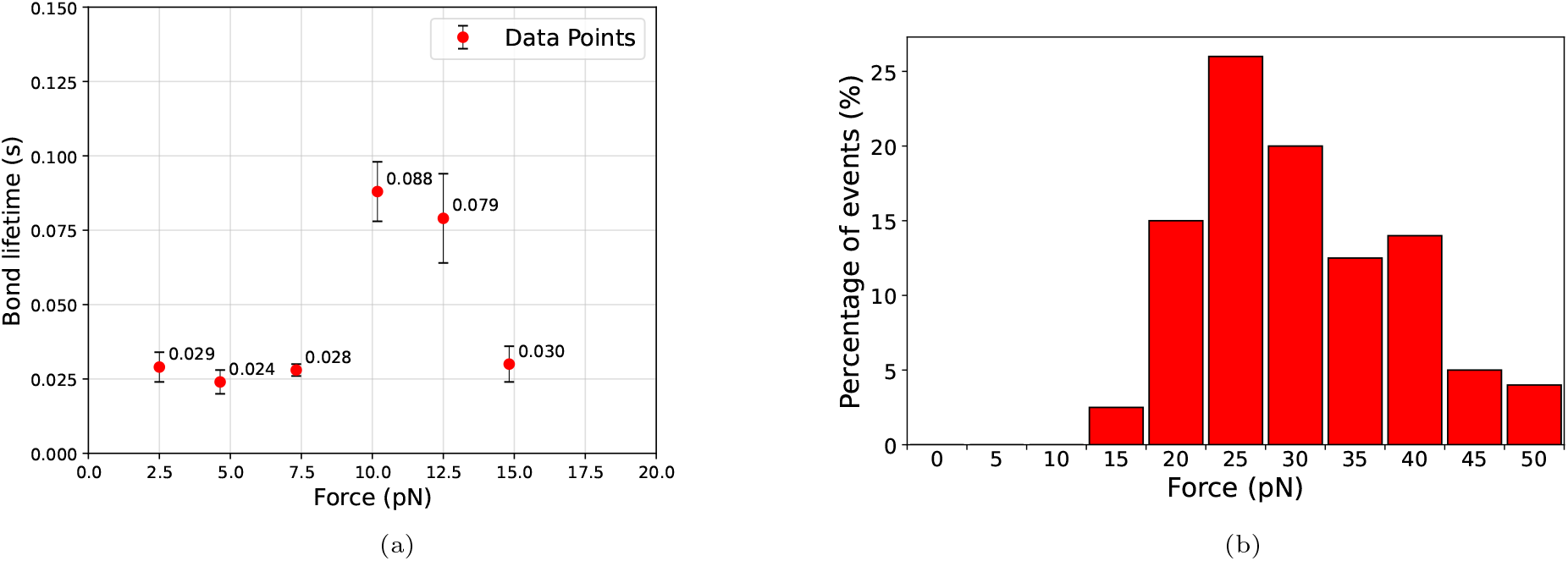
(**a**) Force–lifetime measurements for cadherin bonds, rescaled to N-cadherin. Data points denote mean lifetimes with error bars; values for E-cadherin are taken from [60]. (**b**) Histogram of rupture events of N-cadherin bonds as a function of the applied force. The figure is reproduced using experimental data from [62].

Similarly, studies on E-cadherin have also shown a unimodal right-skewed distribution of rupture events as a function of pulling force. Here, the peak was at 70pN [58]. Interestingly, the right tail was slightly heavier for E-cadherin than for N-cadherin. This is likely due to differences in reproach velocity of the cantilever in the experiments (5*µ*m/s for the N-cadherin vs 3*µ*m/s for the E-cadherin). Taking this into consideration, we assert that the shapes of distributions exhibit a considerable degree of similarity.

Motivated by this similarity, and by the fact that N- and E-cadherin are classical cadherins with homologous binding interfaces, [63], we assume that their force-dependent rupture behaviour has the same functional form/shape and differs primarily by a rescaling of force. Using the ratio of the peak rupture forces, 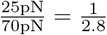 we assume that a force *F* applied to an E-cadherin bond produces the same rupture probability as a force *F/*2.8 applied to an N-cadherin bond. This scaling constitutes a first approximation in the absence of direct N-cadherin force-lifetime measurements. A graphical representation of the resulting N-cadherin force-lifetime data is shown in Figure 2a.

Having established this force rescaling, we next construct models for the lifetimes of catch and slip bonds separately. Because the time of transition between these bond types depends on both force and elapsed contact time, an explicit transition model will be introduced later (Section 3.3). We will start with catch bonds.

### 3.1. Catch Bonds

For N-cadherin catch bonds, we will use the data shown in 2a. However, to account exclusively for catch bond behaviour, we shall consider only the first 4 data points (before the switch from X-dimers to strand-swap dimers). We will first derive an expression for the mean lifetimes of catch bonds, then their *standard deviation* (SD), and finally fit a probability distribution for bond lifetimes.

#### Mean Lifetimes

We extract the lifetime of N-cadherin bonds by using the above-identified factor of 2.8 to account for the relative strength of N-cadherin bonds vs E-cadherin. After extrapolation, we note the change in behaviour around the data point at *F*= 10.18pN.

For biological realism, we want the mean catch-bond lifetimes, *L*_*c*_(*F*), to remain strictly positive for all pulling forces *F* ≥0, and to exhibit approximately constant lifetime at low forces followed by a sharp increase once the force approaches ≈7.5 pN. In [60], the force-lifetime relationship was modelled using a sum of two exponential functions, one for slip bonds and one for catch bonds. We only need to model catch bonds, so we shall use a single exponential function. Consider this standard parametrisation:

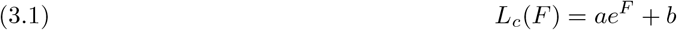

where *L*_*c*_ denotes mean catch bond lifetime in seconds, *F* denotes force in pN, and *a, b* are free parameters. We set *a, b >* 0 to satisfy the criteria outlined above. The *least squares regression* (LSR) (code as Supplementary Material, see Appendix A) yields the following parameter values:

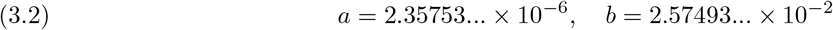

Rounding the parameters to 3 significant digits gives:

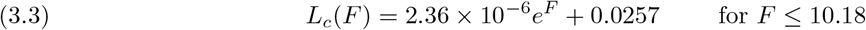

Figure 3 shows the mean catch bond lifetime defined in (3.3).

**Fig. 3:**
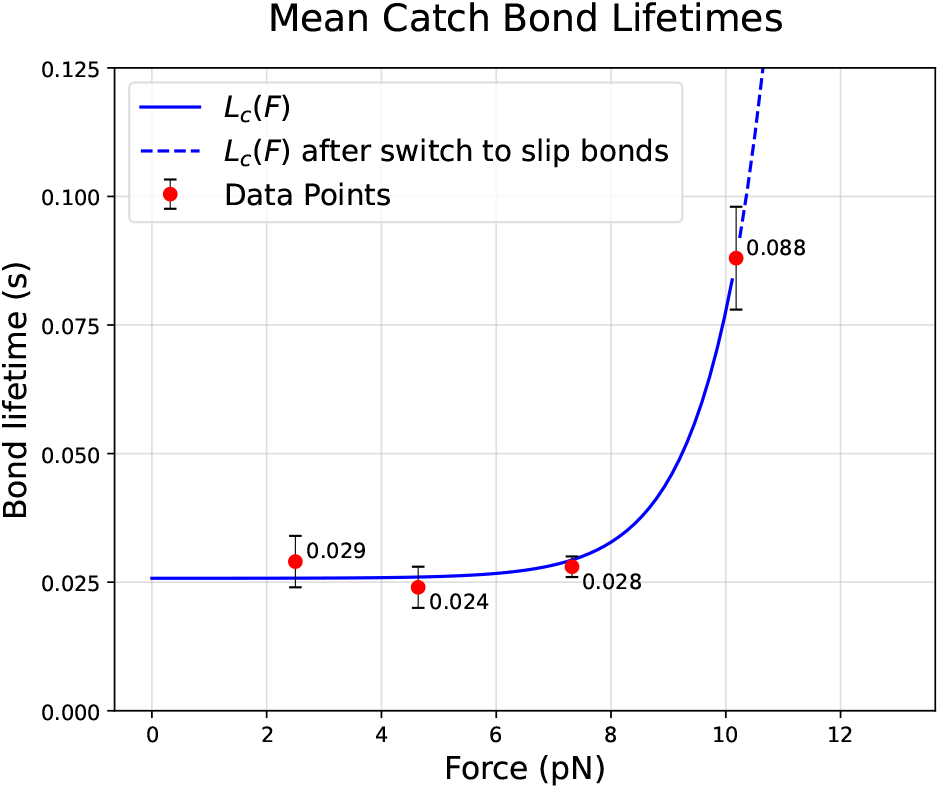
Mean lifetime of catch bonds as a function of the applied pulling force *F*. Red markers denote experimental data points with error bars, while the solid curve shows the fitted catch bond lifetime *L*_*c*_(*F*). The dashed extension indicates the predicted behaviour beyond the catch-slip transition.

#### Distribution of Lifetimes

Having obtained the mean lifetime, *L*_*c*_(*F*), we now derive an expression for the standard deviation of bond lifetimes, *σ*_*c*_. Drawing from [60], we have the data points shown in Table 1.

**Table 1:** Mean lifetimes and standard deviations of lifetimes at given pulling forces, scaled for N-cadherin [60].

| Force (pN) | Lifetime (s) | SD (s) |
| --- | --- | --- |
| 2.50 | 0.029 | 0.027 |
| 4.64 | 0.024 | 0.025 |
| 7.32 | 0.028 | 0.028 |
| 10.18 | 0.088 | 0.061 |

The data suggest that the standard deviation seems to behave similarly to the mean lifetime, in response to growing *F*; we thus assume it is linear,

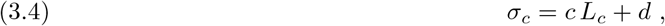

where *a, b >* 0 are constant parameters. Least squares optimization (code as Supplementary Material, see Appendix A) yields the following parameter values:

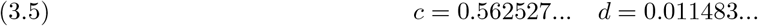

After rounding to 3 significant digits, we obtain the function:

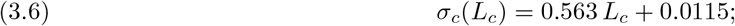

as shown in Figure 4, this produces a very good match for the data points.

**Fig. 4:**
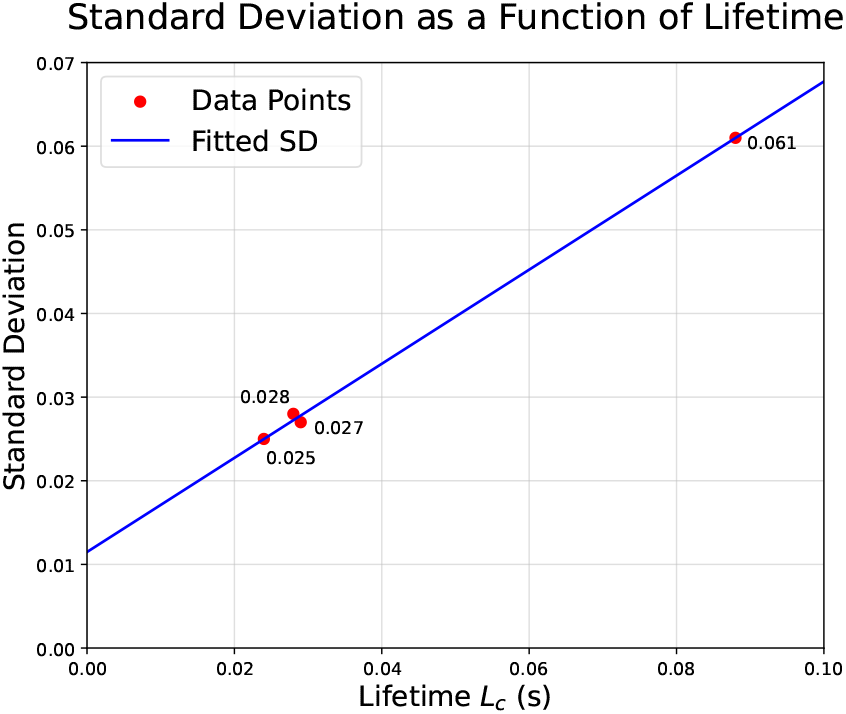
Standard deviation of N-cadherin catch bond lifetimes as a function of the mean lifetime *L*_*c*_. Red markers indicate experimental data points, while the solid line shows the fitted relationship used in the model;cf. (3.6).

Re-writing now (3.6) in terms of the pulling force, using (3.3), we get

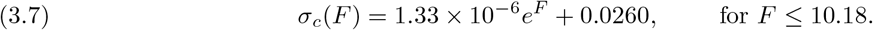

To obtain a full lifetime distribution, we model catch-bond lifetimes using a Gamma distribution, *X*_*c*_ ~Γ(*α, θ*), which provides a unimodal distribution with support on [0, ∞). The parameters are chosen so that the model reproduces the fitted mean and standard deviation:

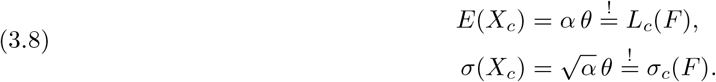

We then obtain the shape and scale parameters of the Gamma Distribution from (3.8), using (3.3) and (3.7):

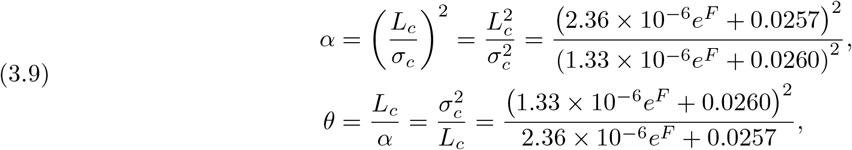

Figure 5 shows the *probability density function* (PDF) and the *cumulative distribution function* (CDF). There the same behaviour can be seen as in the experimental data. Namely, lifetimes are almost independent of the force for *F* = 2.5, 4.64, 7.32 pN (note that the blue *F*=2.5pN line coincides largely with the orange *F*=4.64pN line), followed by a distinct change at *F*= 10.18pN. Note the cluttering around *L*_*c*_ ≈ 0.025s for the first three lines, which coincides with the location of the data points in the experiment.

**Fig. 5:**
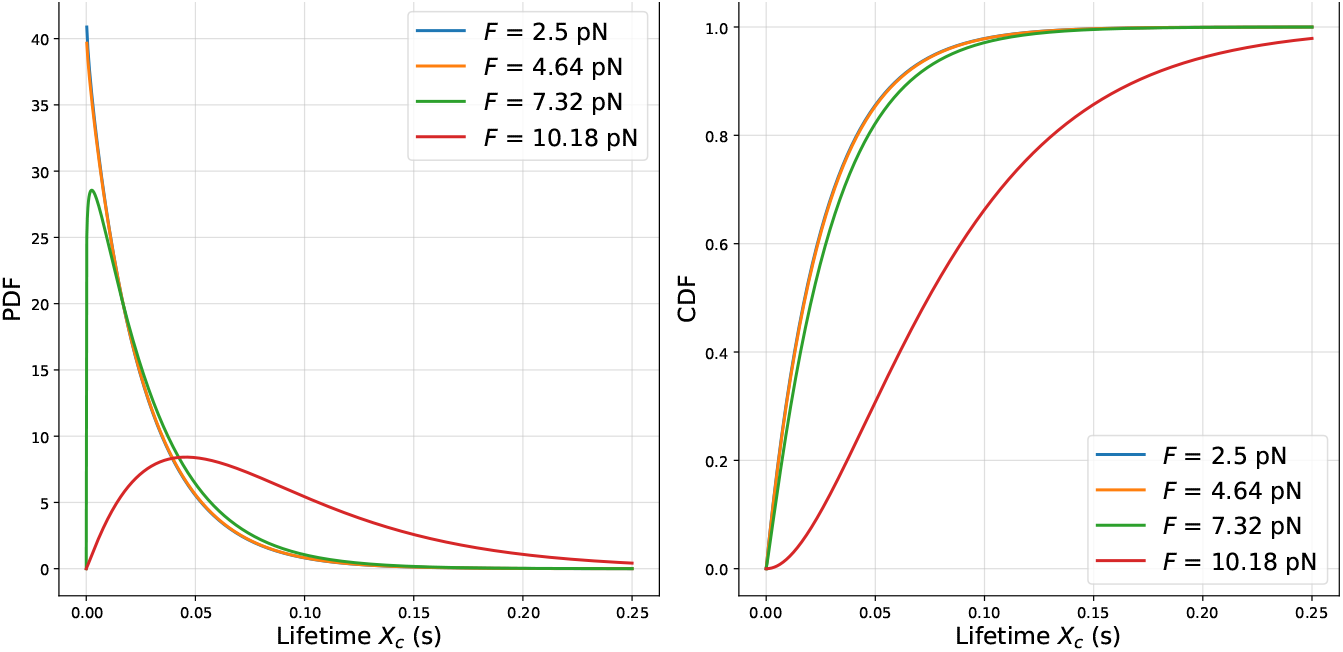
PDF (left) and CDF (right) for the lifetime of catch bonds *X*_*c*_ at the different force values shown in Figure 2a.

### 3.2. Slip Bonds

Moving now to the slip bonds, where in the experiments performed in [60], E-cadherin molecules were forced into their strand-swap conformation (exhibiting slip bond behaviour). Notably, longer contact time before application of a pulling force increased bond lifetime at weaker forces, but did not change behaviour at higher forces. The corresponding N-cadherin data can be found in Figure 6.

**Fig. 6:**
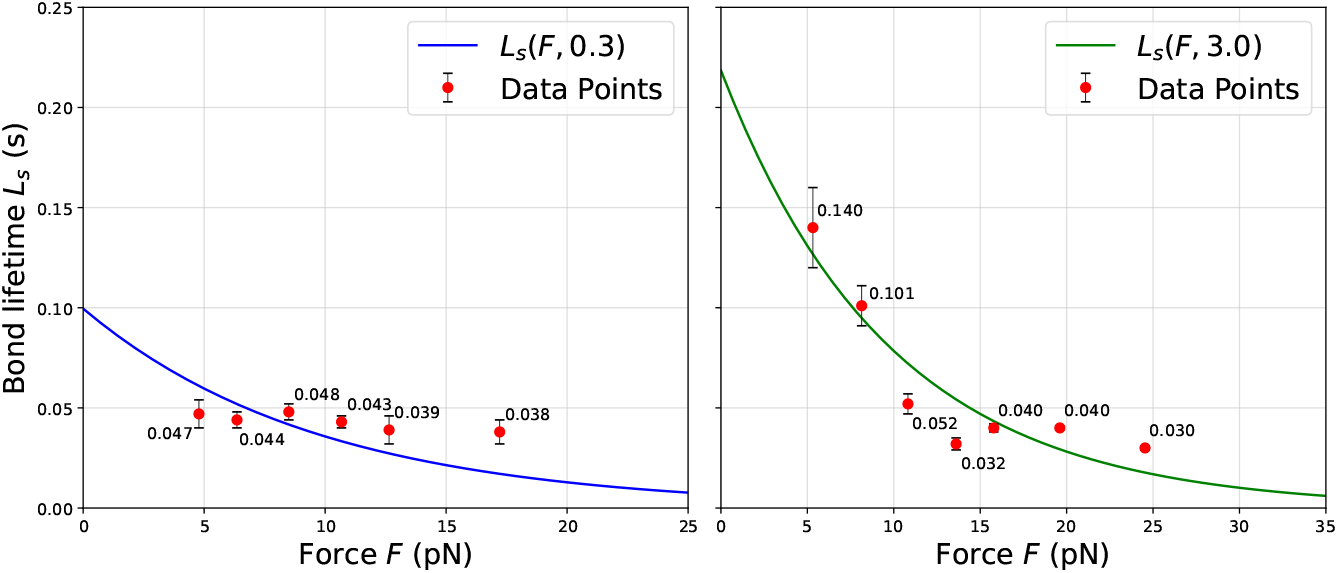
Mean bond lifetime *L*_*s*_(*F, τ*) as a function of the force *F* for contact time *τ* = 0.3*s* (left) and *τ* = 3.0*s* (right). Data points scaled to N-cadherin, data for E-cadherin obtained from [60].

Unlike catch bonds, the experimental measurements depend on a second variable, the contact time *τ* before the application of the force. Consequently, the mean lifetime has to be modelled as a function of both force and contact time.

#### Mean Lifetimes

We model the mean lifetime of N-cadherin slip bonds as a function *L*_*s*_ ≡ *L*_*s*_(*F, τ*) where *F* ≥0 denotes the pulling force (pN) and *τ >* 0 denotes the contact time (s) of the two cells before said force was applied.

Experimental observations indicate that slip-bond lifetimes decrease with increasing force, while longer contact times increase lifetime at low forces and eventually saturate. We therefore assume that bond lifetime decays exponentially with increasing pulling force, and grows in a logistic-type with longer contact time.

To capture the saturation of lifetime with respect to contact time, we model the dependence on *τ* using a logistic equation for *L*_*s*_(*F, τ*):

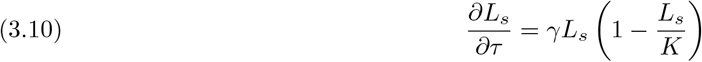

where *γ* is the (constant) growth rate of *L*_*s*_ in *τ*, and *K* = *K*(*F*) is the carrying capacity, the level at which *L*_*s*_ stagnates (which decreases for stronger *F*). After some algebra and integrating (3.10) over time, we have:

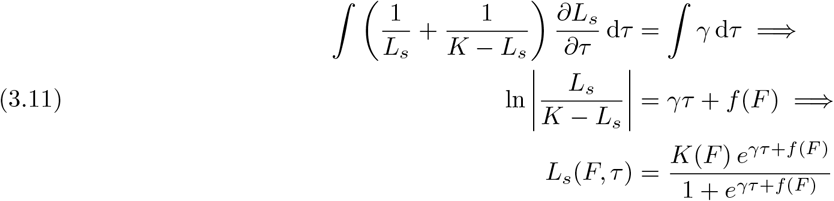

where *f* (*F*) is an arbitrary function of *F*, and *L*_*s*_(*F, t*) takes the form of a sigmoid function. Analogously to our previous assumption about bond lifetimes decreasing exponentially with growing *F*, we now assume that the carrying capacity *K*, which represents the maximum mean lifetime attainable at a given force, decreases exponentially with the applied force *K*(*F*) = *αe*^*−βF*^ where *α, β >* 0 are parameters to be determined. Choosing the simplest possible function for *f* (*F*), namely *f* (*F*) = *δ* = const., (3.11) becomes

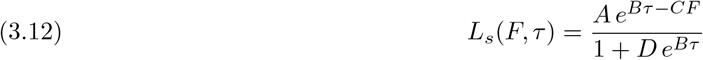

where *A* = *αe*^*δ*^, *B* = *γ, C* = *β, D* = *e*^*δ*^ *>* 0 are constants. Parameter fitting (code as Supplementary Material, see Appendix A) yields:

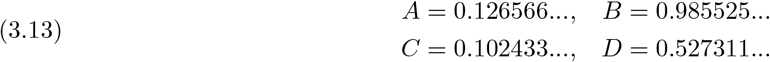

Substituting these values into (3.12) yields the following expression for the mean lifetime of N-cadherin slip bonds (parameters rounded to three significant figures):

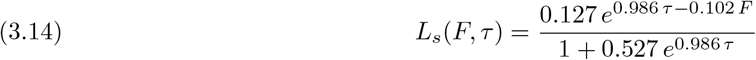

Figure 6, shows that the fitted mean lifetime function *L*_*S*_ for the two different contact times produce a reasonable fit.

The fitted model satisfies the expected qualitative behaviour: lifetimes *L*_*s*_(*F, τ*) remain positive for all *F* ≥ 0 and *τ >* 0, decrease monotonically with increasing force, increase with contact time but saturate, remain finite for all parameter values, and vanish as *F* → ∞.

#### Distribution of Lifetimes

Figure 6 indicates that the standard deviation of slip-bond lifetimes, denoted *σ*_*s*_, is independent of contact times but increases with the mean lifetime, as shown in Figure 7. We therefore assume that contact time influences *σ*_*s*_ only indirectly through the mean lifetime and combine the measurements from both contact-time subgroups into a single dataset, Table 2.

**Fig. 7:**
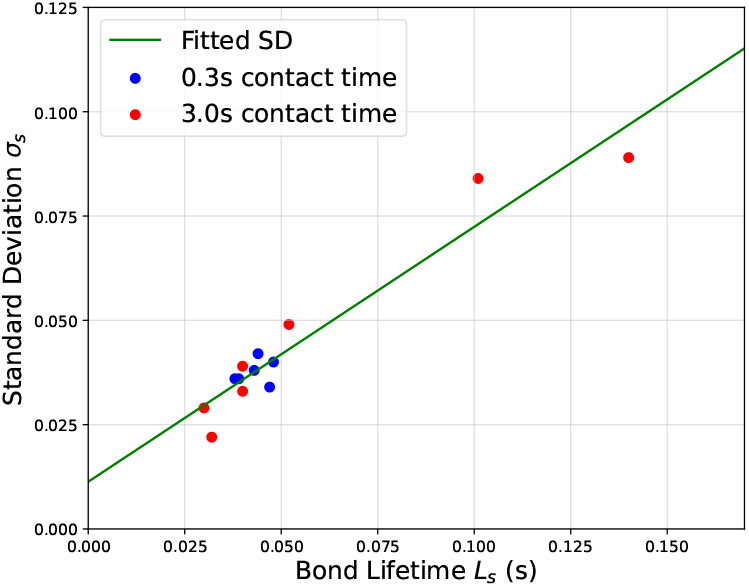
Standard deviation *σ*_*s*_ of slip bond lifetimes as a function of the mean bond lifetime *L*_*s*_. The markers correspond to different contact times, while the solid line shows the fitted (linear) relationship (3.15).

**Table 2:** Pooled pairs of mean lifetime and SD for slip bonds, obtained by combining the data from the two contact time subgroups.

| Lifetime (s) | SD (s) |
| --- | --- |
| 0.140 | 0.089 |
| 0.101 | 0.084 |
| 0.052 | 0.049 |
| 0.048 | 0.040 |
| 0.047 | 0.034 |
| 0.044 | 0.042 |
| 0.043 | 0.038 |
| 0.040 | 0.033 |
| 0.040 | 0.039 |
| 0.039 | 0.036 |
| 0.038 | 0.036 |
| 0.032 | 0.022 |
| 0.030 | 0.029 |

**Table 3:** Parameter values used in the invasion experiment. All spatial quantities are reported in physical units unless otherwise stated. The drift strength *µ*_str_, and the ECM gradient ∇*v* are combined to the drift term of the SDE (4.11) as *µ* = *µ*_str_∇*v*.

| Symbol | Description | Value | Units |
| --- | --- | --- | --- |
| $N$ | number of cells | 250 | |
| $n_c$ | number of catch bonds per cell | 400 | |
| $n_s$ | number of slip bonds per cell | 400 | |
| $R_0$ | initial disc radius | 1.15 | $\mu\text{m}$ |
| $\mu_{\text{str}}$ | drift strength | 0.25 | $\text{s}^{-1}$ |
| $\nabla v$ | (constant) ECM gradient direction | $(1, 0)^T$ | |
| $\lambda_0$ | baseline jump rate | 6.0 | $\text{s}^{-1}$ |
| $R_{\text{int}}$ | interaction radius | 0.345 | $\mu\text{m}$ |
| $F_{\text{switch}}$ | catch-slip bond transition force | 10.18 | pN |
| $n_{\text{rep}}$ | number of stochastic realisations | 10 | |

We can see in Figure 7, that the data points from both experiments (0.3s contact time and 3.0s contact time) follow the same trend, and the shape of the data suggests a linear relationship, so we assume

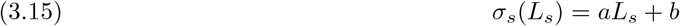

where *a, b* are parameters. LSR (code as Supplementary Material, see Appendix A) yields the following values for *a* and *b*:

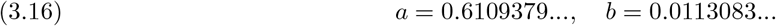

The fit is shown in Figure 7, and now the standard deviation (3.15) reads as follows (parameters rounded to 3 significant digits):

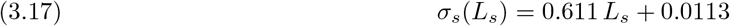

Rewriting now (3.17) in terms of the force *F* and contact time *τ*, using (3.14), we get:

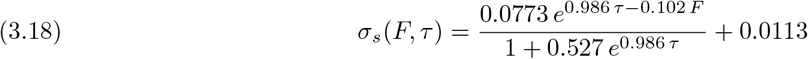

Again, we use a Gamma distribution Γ(*α*^*′*^, *θ*^*′*^) to model the distribution for the lifetime of the slip bonds, for its unimodality and support on [0, ∞). So, for a random variable *X*_*s*_ *~* Γ(*α*^*′*^, *θ*^*′*^), the mean and standard deviation of *X*_*s*_ are given by

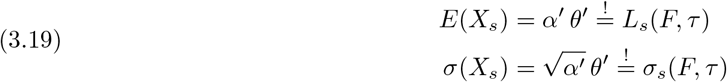

In a similar way as for the catch bonds, (3.14), (3.18), and (3.19) yield

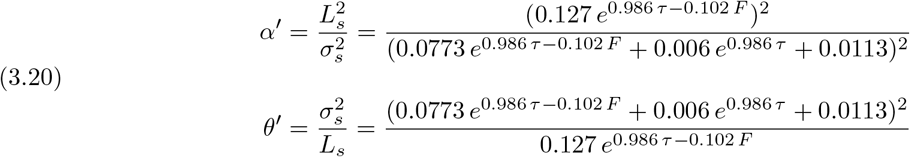

Figure 8, shows the PDF and CDF for *τ* = 0.3s, of the slip bonds lifetime *X*_*s*_, for several values of the force *F* shown in Figure 6. We see the same behaviour as in the experimental data, namely a very weak dependence on force, and very low probabilities to obtain lifetimes above 0.1s.

**Fig. 8:**
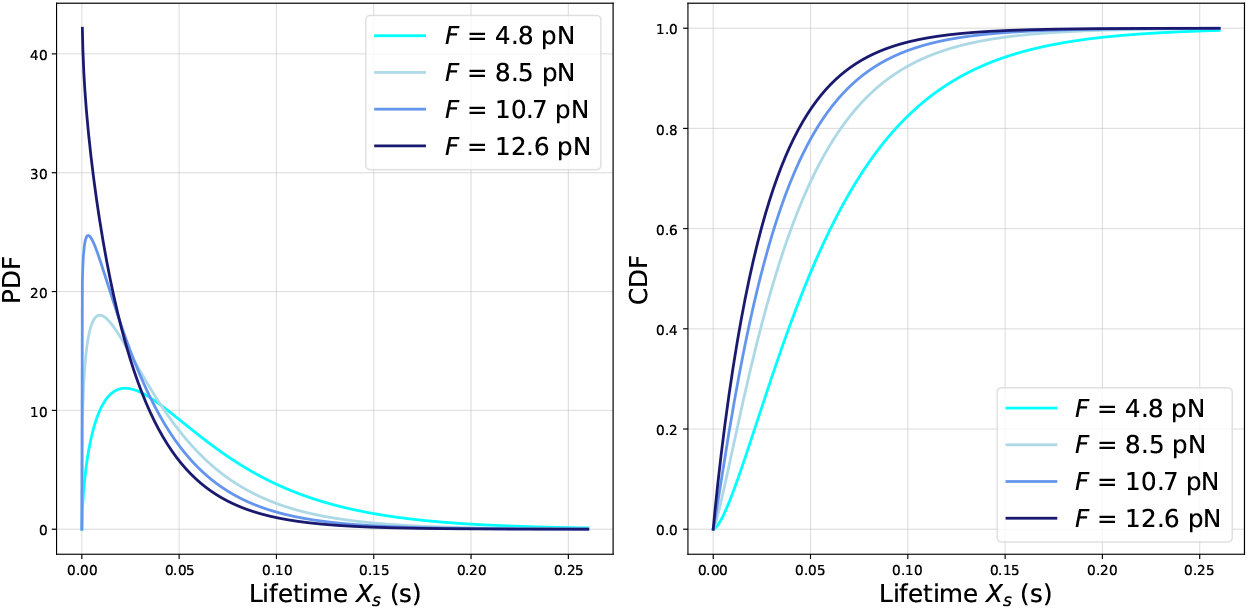
PDF (left) and CDF (right) of slip bond lifetimes *X*_*s*_ for a fixed contact time *τ* = 0.3s and different values of the applied force *F*.

Similarly we plot the PDF and CDF for the slip bond lifetime *X*_*s*_ for *τ* = 3.0s, in Figure 9. Note the slightly higher but still very low probabilities for lifetimes above 0.2s. The behaviour of these functions coincides qualitatively with the experimental data seen in [60]. So, we now go on to consider the conformational switch of the N-cadherin molecule which causes the existence of slip bonds in the first place.

**Fig. 9:**
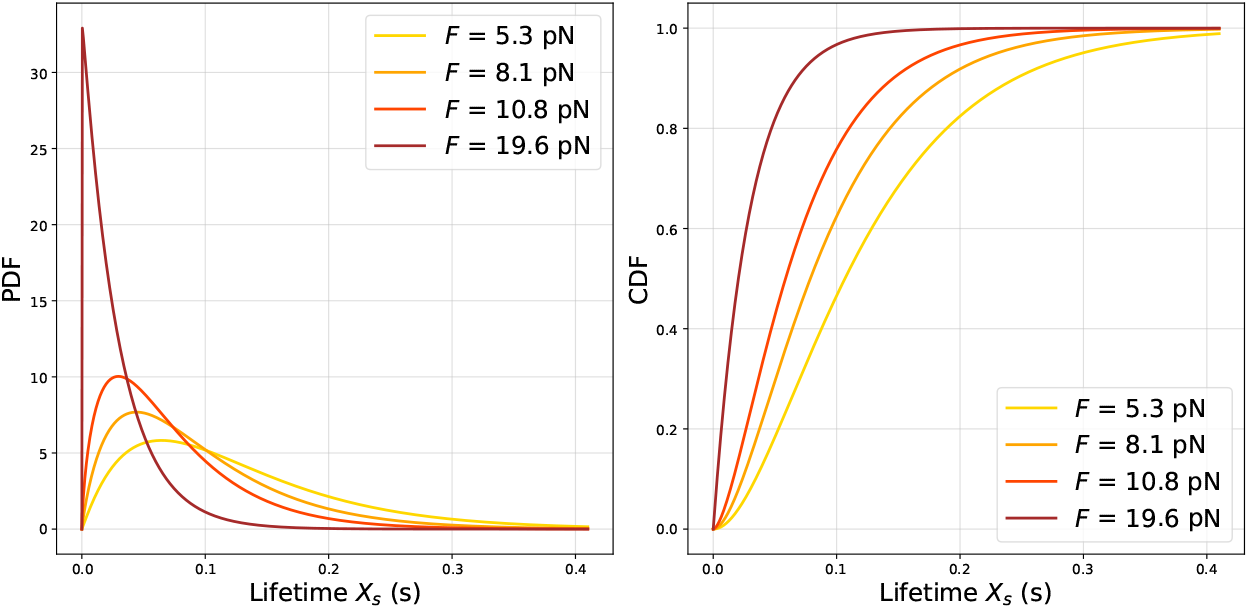
PDF (left) and CDF (right) of slip bond lifetimes *X*_*s*_ for a fixed contact time *τ* = 3.0s and different values of the applied force *F*.

### 3.3. Switch from Catch Bonds to Slip Bonds

Cadherin bonds initially form X-dimers exhibiting catch-bond behaviour and subsequently transition to strand-swap dimers exhibiting slip-bond behaviour. Based on the rescaled data, cf. Figures 2a–3, we assume that all bonds have transitioned by *F* = 10.18 pN and that, in the absence of force, all bonds transition within 1 s of contact. The transition is assumed to be instantaneous.

Now we want to determine the time required until the N-cadherin molecule undergoes this transformation. Define 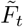 as the pulling force acting on the cell at time *t*. For model simplicity, we assume that this force can be treated as a constant over the time periods in question. Furthermore, define 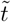 as the time elapsed since bond formation.

We define the cumulative probability (in the sense of a cdf) 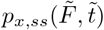 that a bond transitions at or before time 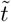 under force 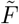. We choose a simple model with linear dependence on force and time since bond formation,

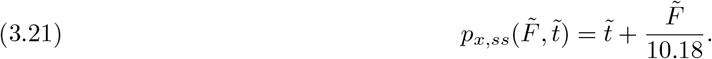

This choice reflects the assumptions that all bonds have transitioned at 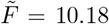 and, in the absence of force, by 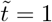. Note that, since we assume 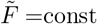., *p*_*x,ss*_ grows linearly with time.

Now define *U ~* Uniform[0, 1]. The bond will transition when 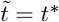 such that

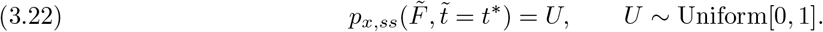

Thus, *t*^∗^ represents the time between bond formation and transition from catch bond to slip bond.

Then, using (3.21), we can express (3.22) as a function of the transition time:

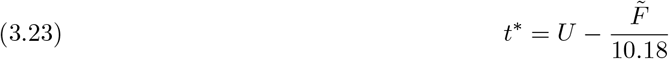

This expression returns the time after bond formation at which the transition from catch bond to slip bond occurs. If the returned value is negative, we take *t*^∗^ = 0, i.e. we assume instantaneous transition of the bond.

If the pulling force is applied more than 1 s after bond formation, all bonds are assumed to be in the strand-swap (slip-bond) regime, and the contact time *τ* is measured from the transition.

If the pulling force is applied less than 1s after initial contact, either the catch bond ruptures before the transition or the transition occurs first. In the latter case, the slip-bond model applies with *τ* = 0, giving

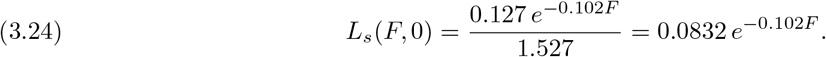

Now that we have derived expressions for the lifetimes of N-cadherin catch and slip bonds, and their conversion process, we can turn our attention to the mechanism of how these quantitative metrics of adhesion can alter the diffusion term of an existing model.

## 4. Incorporating Cell-Cell Adhesion into the Diffusion Term

In this section we incorporate the effects of cell-cell adhesion into our MCC individual based model (2.1), and in particular into the diffusive component 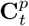, cf. (2.2), of the SDE.

Cell-cell adhesion mechanically stabilises neighbouring cells and therefore reduces their random motility. In contrast, the drift term is driven by chemotactic and haptotactic cues and is assumed to remain unaffected. Therefore, adhesion acts by reducing the stochastic component of motion governed by the compound Poisson process 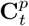. This leads to more coordinated, directed migration along chemical and mechanical gradients, rather than the more pronounced random motility exhibited by single cells.

As described in Section 2, 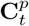 models the timing and number of random jumps through a Poisson process *𝒩*(*t*), as well as the direction and magnitude of the random jumps through a collection of i.i.d. random variables 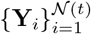.

Following the assumptions made above, we will incorporate cell-cell adhesion dynamics by dereasing

- the frequency of jumps, governed by *𝒩*(*t*),
- the magnitude of jumps, governed by the magnitude of the 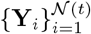.

We address each one separately in the following sections.

### 4.1. Decreasing the frequency of random jumps

Adhesive bonds between two MCCs stabilise both of them, and partly block pseudopodia sensing in the direction of the other cell. This leads to a lower turning frequency for both cells. To quantify this, we estimate how much of the surface area of one cell is obstructed by another nearby cell.

We define the degree of obstruction of cell *p*_*k*_ by *p*_*j*_ as *γ*_*k,j*_ by the relative overlap of the cell surfaces,

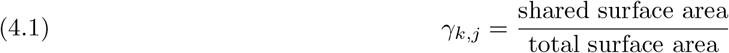

Based on this expression, we can determine the new jump rate 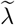 of (*t*), by introducing the total obstruction as the sum of obstructions caused by nearby cells:

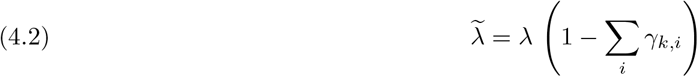

where the *p*_*i*_ are cells within the interaction radius of *p*_*k*_. One consequence of this expression is that cells that are completely surrounded by other cells, i.e.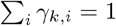, tend to exhibit zero random motility, hence their migration becomes deterministic. In other words, random movement is limited to cells that are found at (or close to) the boundary of the body of cells. A geometric example of this obstruction model can be found in Appendix B.

### 4.2. Decreasing the magnitude of random jumps

The random motility of MCCs decreases with the creation of adhesive bonds. In particular, stronger and longer-lasting bonds result in greater mechanical stabilization. We thus want to scale the jump magnitude by a factor inversely proportional to the lifetimes of the cell bonds.

We define the sum of lifetimes of bonds between cells *p*_*k*_ and *p*_*a*_, *α*_*k,a*_, by:

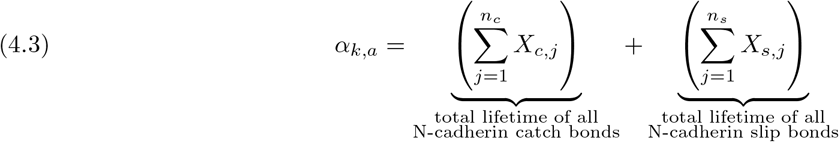

where *n*_*c*_, *n*_*s*_ denote the number of catch bonds and slip bonds, and *X*_*c,j*_, *X*_*s,j*_ denote their respective lifetimes. Note that we assume the *X*_*c,j*_, *X*_*s,j*_ to all be independent and, within their specific bond type, identically distributed. Therefore, if the number of cell bonds is large enough, we can invoke the Central Limit Theorem (CLT), which states that the mean of the *X*_*i*_ (where *i* = *c, s*), which we denote by 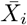, is a random variable approximately following a normal distribution with mean 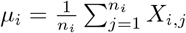, and variance 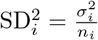. Here, *σ*_*i*_ is the standard deviation of lifetimes of bond type *i* that we found earlier, and note that we defined the *X*_*i,j*_ based on their mean, *L*_*i*_. Therefore,

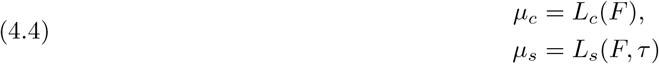

and so,

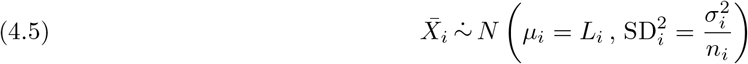

where “ 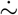“ represents the “follows approximately” symbol. We can now use the random variable 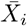 as an expression for our bond lifetimes. Equation (4.3) will then equate to

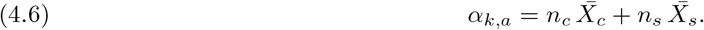

We now introduce a reduction factor *f* to scale the jump magnitude in an inversely proportional manner to the lifetimes of the cell bonds; for a pair of cells *p*_*k*_ and *p*_*a*_, we define

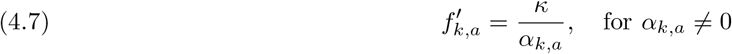

where we included a multiplicative constant *κ* to keep the system adaptive to match experimental data. *κ* has unit [*n*_*k*_]*·*[*L*_*k*_] (the unit of the number of bonds times the unit of lifetimes) so as to make *f* ^*′*^ dimensionless. We want to exclude situations where there are no bonds between two cells, i.e. *α*_*k,a*_ = 0, thus we sum over all cells *p*_*i*_ within the interaction radius of *p*_*k*_, and introduce indicator variables *D*_*k,i*_ to specify which cells have bonds between them. In particular, consider cell *p*_*k*_ with cells *p*_*i*_, *i* = 1, 2, 3,*…* in its vicinity. Then, we define the reduction factor *f*_*k*_ for *p*_*k*_ as

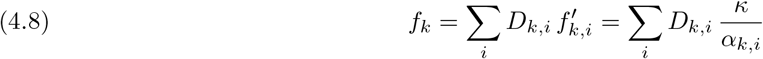

where the indicator variable *D*_*k,i*_ is defined as

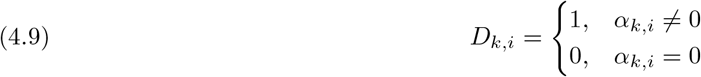

Hence, for large enough numbers of bonds—in a typical cell-cell interaction scenario, there will be hundreds if not thousands of bonds, so invoking the CLT is justified, despite the asymmetry of the Gamma distribution— we can define our modified CPP jump magnitude 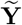-for *p*_*k*_ as:

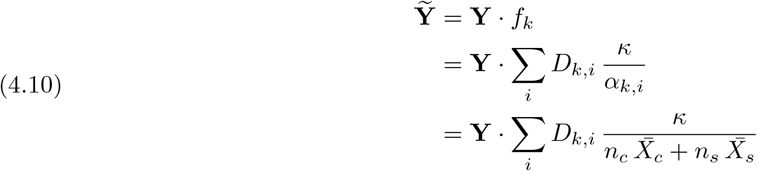

With this formulation, increasing the number of bonds and their lifetimes reduces the magnitude of random jumps. The jump magnitude also depends implicitly on the pulling force: beyond the lifetime peak, stronger forces shorten bond lifetimes, which in turn increases jump magnitude and promotes cell separation. The model therefore captures a key feature of adhesion, namely that long-lived bonds stabilise cell pairs, whereas shorter-lived bonds have a higher likelihood of separation.

### 4.3. The Updated Movement Equation for Mesenchymal Cancer Cells

We now combine the previous results to obtain the updated movement equation for MCCs that reads:

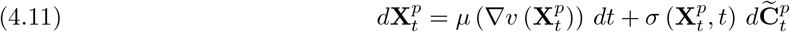

where the updated compound Poisson noise term is given by:

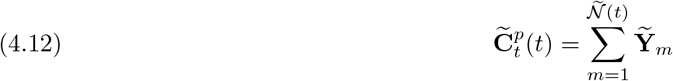

where 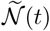 is a Poisson Process with rate 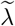 given by (4.1) & (4.2), and 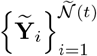 are i.i.d. vector-valued random variables given by (4.5) & (4.10). The newly formed CPP incorporates directly the adhesion dynamics of catch-slip bonds of N-cadherins, since the mean and standard deviation of (4.5) are explicitly given through our parameter estimation on experimental observations, performed in Section 3. In particular, the mean values are given by (3.3)-(3.14), and the standard deviations by (3.7)-(3.18), for catch and slip bonds, respectively.

We show the overall behaviour of the bonds graphically in Figure 10. It illustrates the saturation of slip-bond lifetimes with increasing contact time *τ*, consistent with the logistic growth assumption built into the model. The figure also highlights the two dominant factors governing bond lifetimes: the rate of the transition from catch to slip bonds, and the contact time *τ* in the slip-bond regime. Together, these mechanisms play a central role in determining the overall migratory behaviour of the cells.

**Fig. 10:**
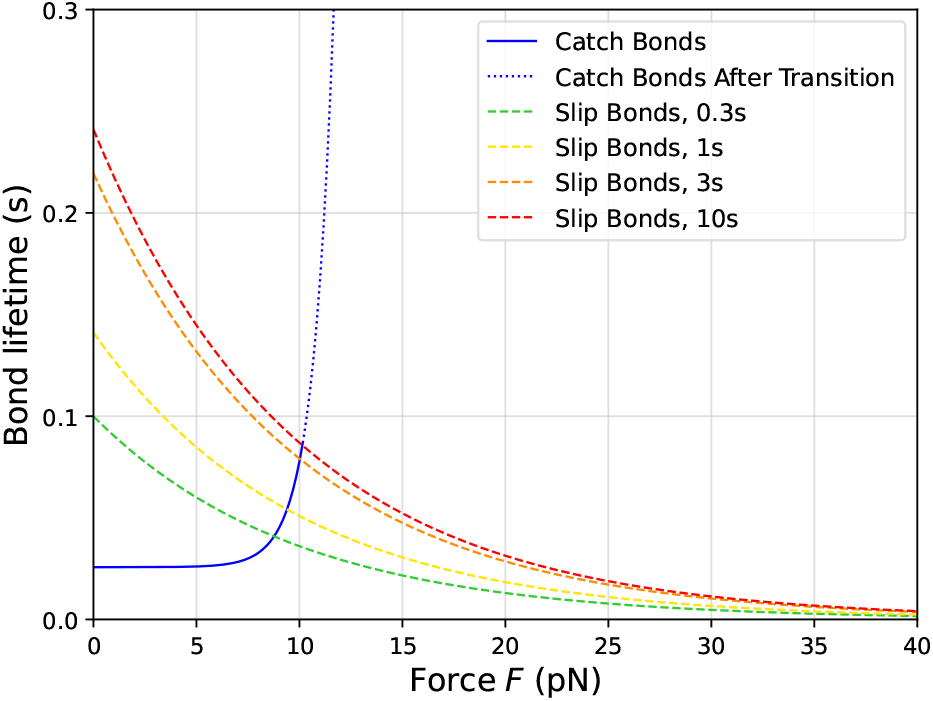
Showing the mean bond lifetimes as functions of the applied force F for catch and slip bonds. The solid blue curve represents the catch bond lifetime before the catch–slip transition, while the dotted blue the continuation beyond the transition. The dashed curves correspond to slip bond lifetimes evaluated at different contact times, as shown in the legend.

## 5. Results

In this section we investigate how force dependent adhesion affects the migratory properties of the population of the updated stochastic cell motility model (4.11)-(4.12) summarised in Section 4.3.

The goal is to deliver a qualitative comparison between the baseline (2.1)-(2.2) and the adhesion (4.11)-(4.12) model. The numerical implementations preserve the structure of the model: a drift term describing the biased part of the migration combined with a compound Poisson process for the random part. Cell-cell adhesion is accounted for by reducing the effective jump rate (through local obstruction, Section 4.1) and the effective jump magnitude (through force dependent bond lifetimes, section 4.2), as described in Section 4.

For the sake of computational efficiency and clarity of presentation, the numerical simulations presented below primarily focus on population level transport effects. Hence, we have employed the following modelling simplifications.

- Two cells are assumed to always be in contact if their distance is below a prescribed radius *R*_int_ *>* 0. Within this radius, it is assumed that the two cells are always in contact. This simplification replaces the stochastic bond formation and rupture by an average static contact. In effect, it removes the need to explicitly track cell-cell contact histories while maintaining the local nature of adhesion.
- The obstruction term *γ*_*k,i*_ employs the shared surface area between the two cells that is approximated through, and decreases with, their interaction distance *R*_int_.
- We approximate the average lifetime across many bonds by a Gaussian distribution. In the numerical simulations, we employ the CLT level of the bond lifetime aggregation, appropriate for the large bond number regime considered. This simplification has a significant impact in the computational cost.
- Transitions between catch and slip phases are represented through force dependent bond lifetime statistics evaluated at a representative contact time, rather than through explicit time dependent switching probabilities.

For the comparison of the baseline model with the the adhesion system, we use various metrics: spatial extent and organisation of the population, and force-dependent motility statistics. These metrics reveal a clear transition from diffusive spreading of the population to a confined regime.

The spatial spread of the population, is characterised/classified through the radius of the gyration

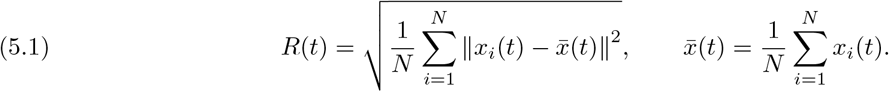

In Figure 12, the spatial spread of two initially densely packed populations of cells was simulated over time. The baseline population (blue) was assumed to exhibit no cell-cell adhesion, while the adhesion population (orange) was initiated according to our regime. The figure clearly shows shows a qualitative difference between the two regimes. In the baseline case, the radius of gyration grows almost monotonically; this is consistent with diffusive spreading. In contrast, the radius in the adhesion model, approaches a plateau and remains almost constant throughout the simulation. We thus conclude that force-dependent adhesion induces a confinement that prevents spreading while maintaining local fluctuations.

### Spatial organisation of the cell population

Figure 11 exhibits the confinement of the adhesion system as opposed to the baseline case. In both cases, the two populations move along a gradient towards the right-side of the domain. The baseline case shows gradual spatial spread of the population, a characteristic of diffusive growth. In contrast, in the adhesion case, the population remains compact, with only small changes in shape and position. These dynamics are consistent with the quantitative measurements: adhesion stabilises the population.

**Fig. 11:**
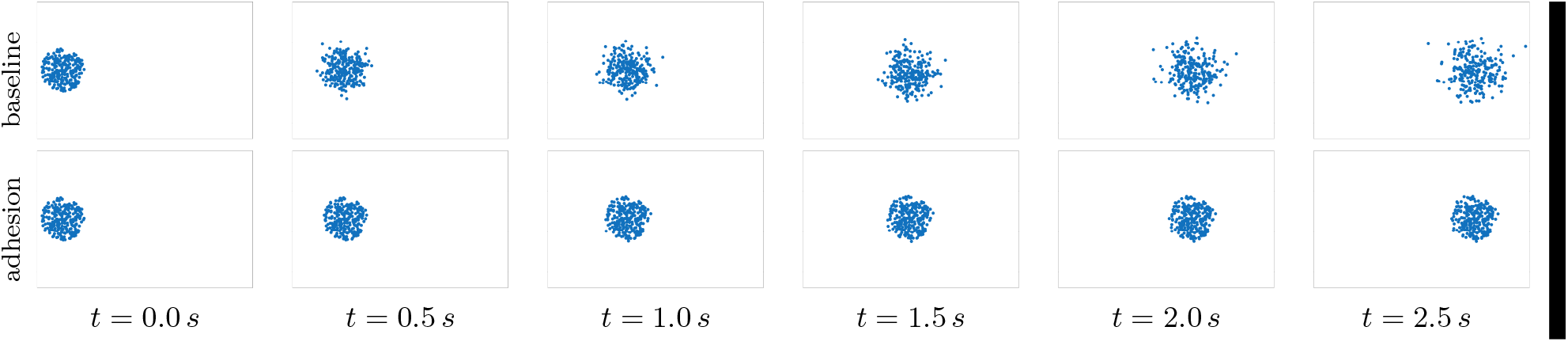
Snapshots of the cell population of 250 cells over the domain [0, 11.5] × [2.3, 9.2] *µ*m^2^ at various time points for the baseline (top row), and the adhesion (bottom row) dynamics. The population in the baseline spreads-out progressively, whereas the adhesion-modified population remains compact over long times; it only exhibits small stochastic fluctuations in shape and position.

**Fig. 12:**
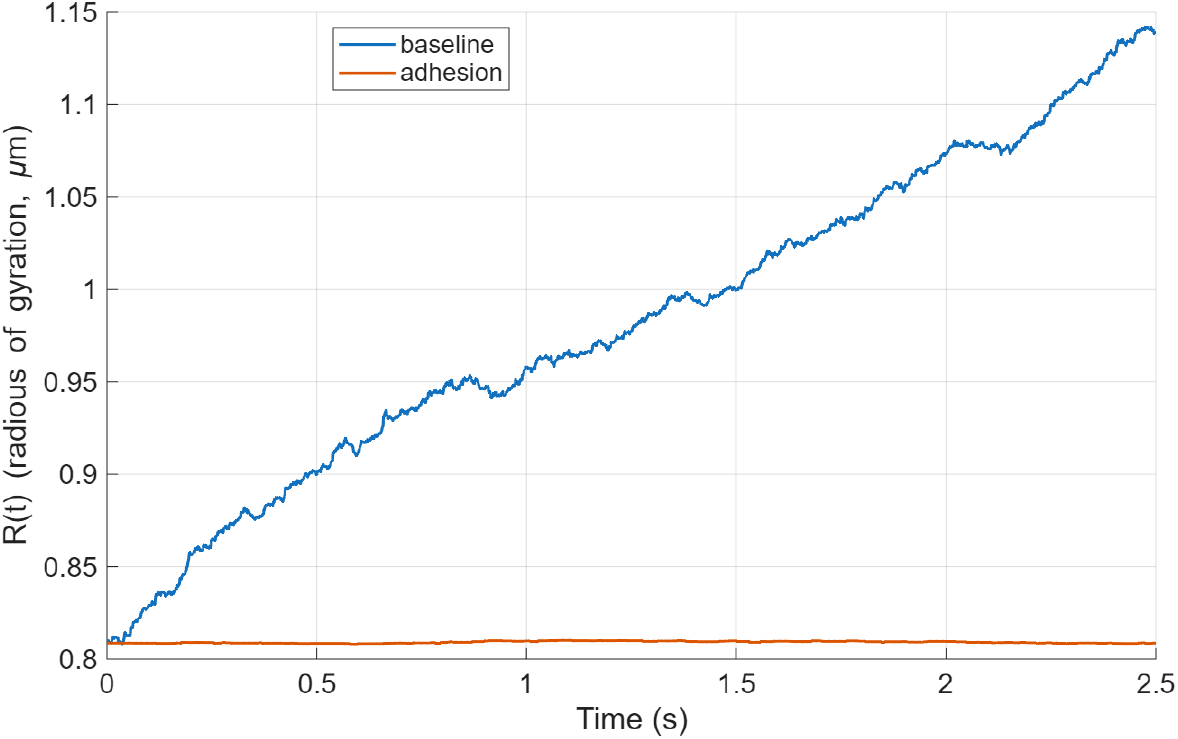
Evolution of the population radius *R*(*t*) in the baseline and adhesion cases. The baseline model exhibits monotone growth of *R*(*t*) consistent with diffusive spreading, whereas the adhesion model rapidly approaches a plateau. This indicates the emergence of a dynamically confined stochastic equilibrium. The clear separation of the two curves indicates that force-dependent adhesion induces confinement rather than merely slowing diffusion.

To further investigate the robustness of the confinement mechanism, we performed an additional numerical experiment using strip initial conditions. The corresponding dynamics are shown in Figure 14. Therein it shows that in the absence of adhesion, the initially narrow strip progressively broadens due to stochastic spreading. In contrast, under force-dependent adhesion, the strip remains coherent, preserving its overall shape. This confirms that the emergence of confinement is not specific to radially symmetric initial data, but persists under other configurations.

Figure 13 shows the *mean squared displacement* (MSD) at final time *T* as a function of the applied force *F*. The adhesion model, incorporates, through the bond lifetime mechanism, force-dependent control of both the jump frequency and jump magnitude. On the other hand, in the absence of adhesion in the baseline case, cell trajectories follow a compound Poisson process with constant rate. The latter, leads to diffusive spreading at the cell population level. Across the force range, the adhesion model exhibits a clear reduction in MSD relative to the baseline dynamics. A clear separation of scales is evident: adhesion reduces the stochastic diffusivity by approximately two orders of magnitude.

**Fig. 13:**
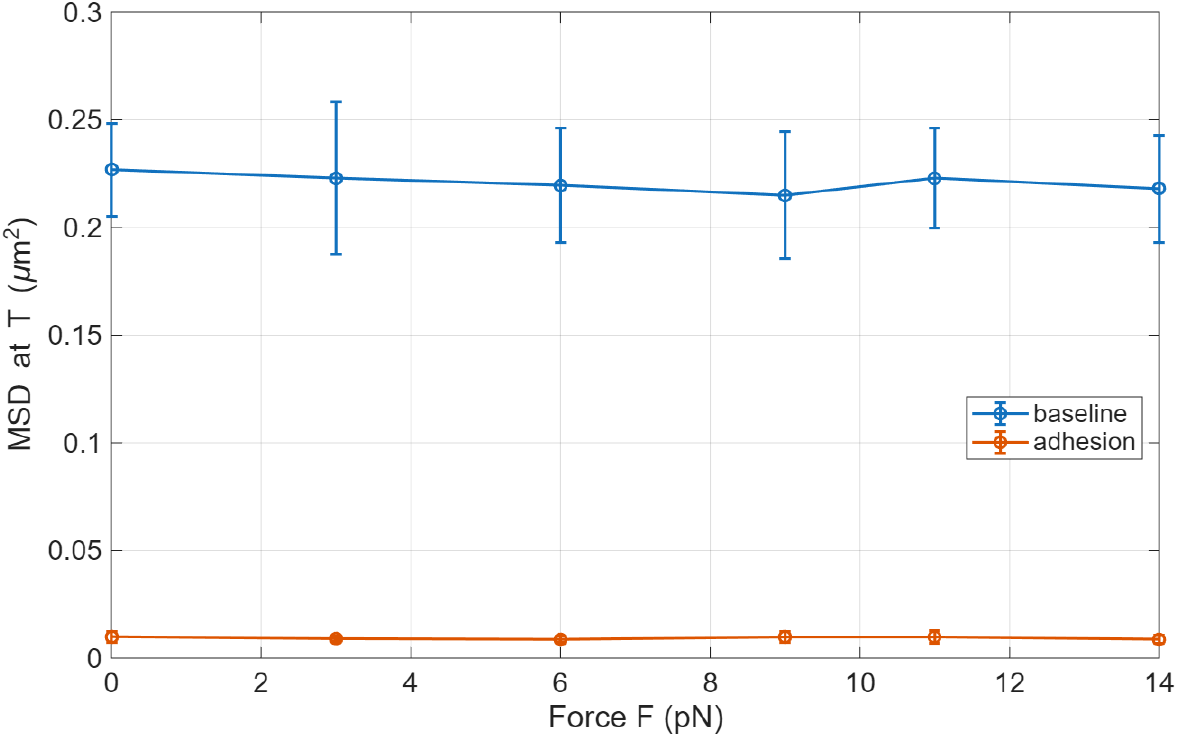
Mean squared displacement (MSD) at final time *T* as a function of applied force *F*. The baseline dynamics produce large, weakly-force dependent MSD consistent with unconstrained diffusive motion. In contrast, the adhesion model reduces significantly the effective MSD across the entire force range.

**Fig. 14:**
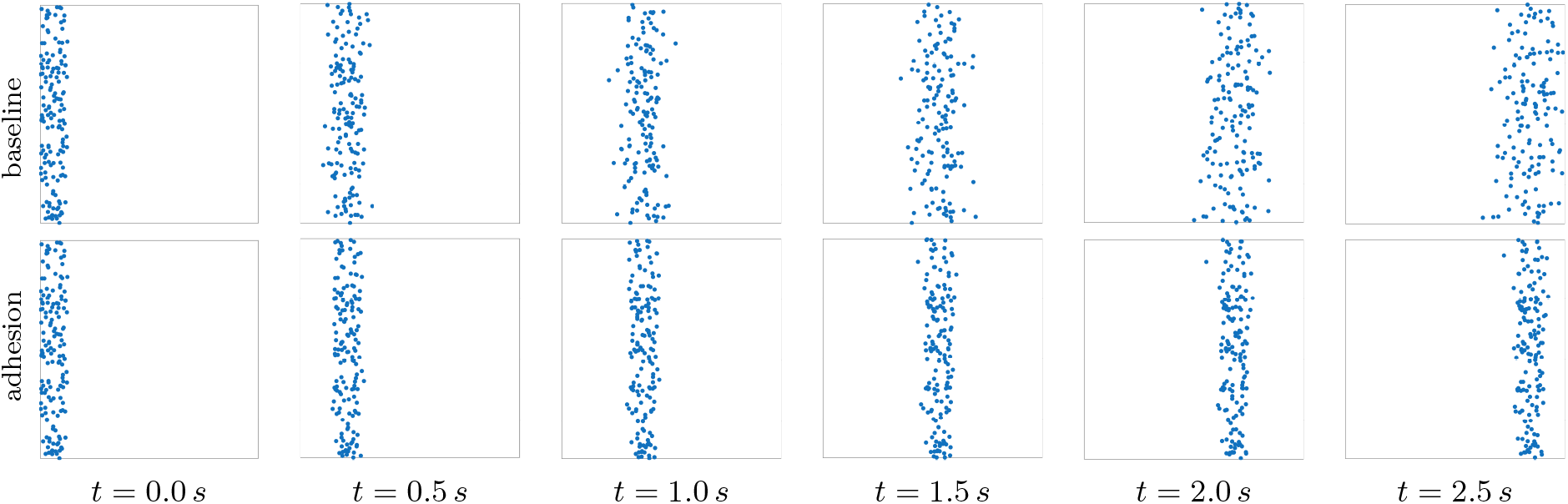
Snapshots of the cell population (250 cells) for the numerical experiment with strip initial conditions over the domain [0, 9.2] × [2.3, 9.2] *µ*m^2^ at times *t* = 0, 0.5, 1.0, 1.5, 2.0, 2.5 s. Top row: baseline dynamics. Bottom row: adhesion-modified dynamics. Clearly, under the baseline model, the initially confined strip progressively spreads-out. In contrast, the adhesion preserves a compact, coherent strip, with only small stochastic fluctuations in shape.

Taken together, the results presented in this section are evidence that force-dependent adhesion introduces a distinct dynamical phase. The system transitions from a diffusive spreading regime to a confined one. This confinement arises naturally from the bond-lifetime and does not require explicit external or artificial boundary forces.

## 6. Conclusion

The goal of this work was to introduce a mathematical modelling framework that allows cell-cell adhesion forces to be incorporated into individual based cell migration models. The driving idea was that the mechanical stabilisation that arises from adhesion bonds, would decrease the random migration of MCCs, without altering their directed migration.

We showed that the lifetime of a single adhesive bond can be expressed as a random variable, for which the expressions for mean and standard deviation were derived from experimental data. Based on these quantities, probability distributions were selected to ensure biological realism and parametrised accordingly.

This stochastic description was subsequently used to modify the diffusion term of the MCC migration model in a force-dependent fashion: increasing number of bonds and longer bond lifetimes lead to a reduction in random motion.

The numerical results show that this mechanism induces a qualitative change in the migration dynamics at the population level. In the absence of adhesion, populations of MCCs exhibit the expected diffusive spreading. In contrast, force-dependent adhesion gives rise to a confinement regime, in which the population remains coherent. This confinement emerges naturally from the bond lifetime mechanism and does not require external forces or boundary mechanisms. The results further indicate that adhesion reduces the stochastic diffusivity by orders of magnitude.

In this respect, the approach presented here is novel, as it targets cell–cell adhesion from a stochastic rather than a deterministic perspective, while still being directly informed by experimental measurements. Moreover, modelling the transition from catch bonds to slip bonds as a dynamic process influenced by both force and contact time provides an alternative to the commonly employed static threshold descriptions.

Although there is a limited availability of biological data, the model is able to capture a well established qualitative phenomenon, namely the stabilising effect of intercellular adhesion on migrating cancer cell populations. This suggests that the framework could serve as a motivation for further experimental studies on force dependent N cadherin adhesion.

An extension of the model would be to incorporate a refractory period following bond rupture and to allow for subsequent rebinding events. Such extensions would enable a more detailed representation of the adhesion dynamics and further influence the motility of the cell population. Furthermore, the addition of volume filling effects through the inclusion of cell-cell repulsion forces into the current model will ensure the physical separation of overlapping interacting cells.

From a modelling perspective, this work constitutes a step towards improving predictions of invasion dynamics, embedding cell-cell adhesion into multiscale invasion models where stochastic effects influence macroscopic dynamics, and potentially informing therapeutic strategies that target adhesive interactions.

## Appendix A. Code availability

All numerical simulations and visualisations presented in this paper were conducted in MATLAB [64] and Julia [65]. The full source code used, is publicly available at: https://github.com/sfaknikj/force-dependent-adhesion-cell-migration

The repository includes a `README` file describing the structure of the code, parameter settings, and instructions for reproducing the computational results. The computational workflow consists of the following steps:

- simulation of cell motility dynamics driven by a drift term and a compound Poisson process;
- implementation of force-dependent adhesion effects through obstruction-modified jump rates and bond-lifetime-modulated jump magnitudes;
- generation of population-level observables, including spatial snapshots, radius of gyration, and force-dependent motility statistics;
- visualisation of baseline and adhesion-modified dynamics for qualitative and quantitative comparison.

### Licence

The source code is distributed under the [MIT Licence](https://opensource.org/license/MIT). This permits reuse, modification, and redistribution of the software under the terms of the licence, provided that the original copyright notice is retained.

Use of this software in academic work should be accompanied by an appropriate citation of the present paper. If the code contributes to published results, users are kindly requested to reference this article as the primary source describing the modelling framework and numerical implementation.

## Appendix B. Example Model of Cell Overlap

As a rough conceptual estimate, we approximate a cell’s shape by a rectangular cuboid. We furthermore assume that all cells are approximately of the same size and shape. Figure 15a shows how two such cells would look next to each other. The amount by which the cells interfere with each other is determined by the ratio of the area of the face obstructed by the other cell, to the total surface area of the cuboid. We shall furthermore assume that the width of the cells is equal to their height, and thus, we only need the ratio of their length to their width.

**Fig. 15:**
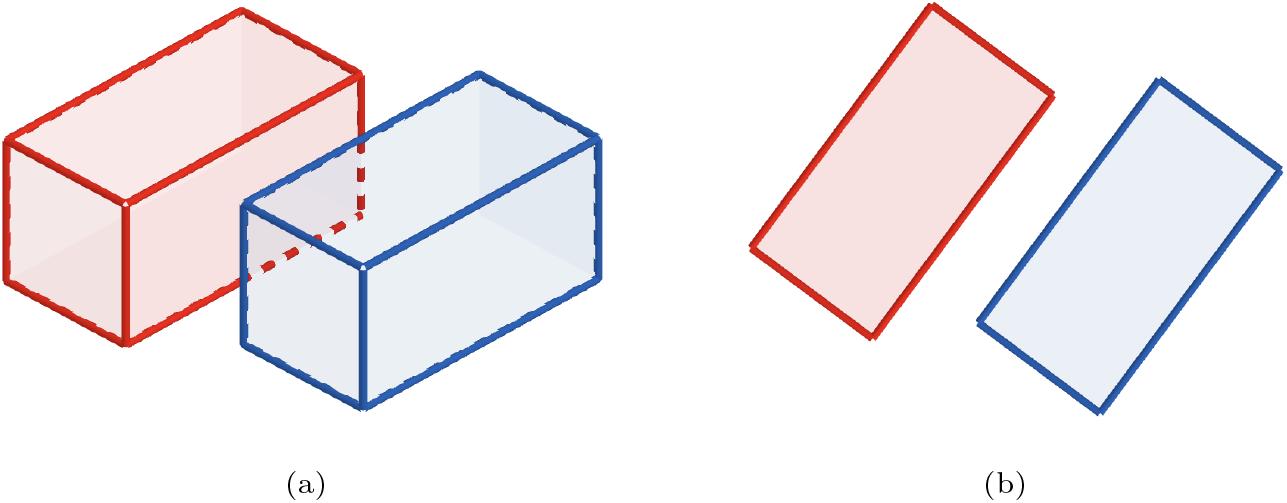
Schematic representation of the two cells modelled as rectangular cuboids overlapping along their long axis. (a) Side view: the dashed region indicates partial overlap/obstruction along the long faces. (b) Top view: relative orientation of the cells and the projected overlap/obstruction.

A study investigating mouse, cat, and dog cancer cells (which we assume to be similar enough to human cancer cells, at least in the context of this basic model) showed a mean inner diameter (assuming cells are elliptical, this will be the major axis) of 11.5µm, with circularities ranging from 0.6 to around 0.95 [66]. The circularity of the elliptic cross section of a cell is given by

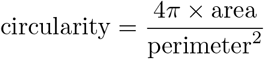

We find most cancer cells to have circularities of close to 0.95, [66], but to account for low outliers, we shall go with a circularity of 0.9. In the case of an ellipse with major axis *a* and minor axis *b*, the area is given by *πab*. Thus,

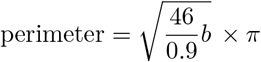

where we used *a* = 11.5µm, as mentioned above. We find *b* ≈ 6.9. Therefore the ratio for the width/height of the cell to its length, is given by 6.9µm: 11.5µm = 0.6: 1. Using the cell length as unit length, the faces in the length-direction have an area of length *×* height (or width) = 1 *×* 0.6 = 0.6. The smaller faces have an area of height *×* width = 0.6 *×* 0.6 = 0.36, and thus we get a total surface area of

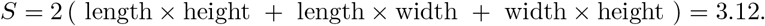

Of course, the cells aren’t always perfectly aligned. We compute *γ*_*k,j*_, the degree of obstruction of cell *p*_*k*_ by *p*_*j*_ by the relative overlap of the cell surfaces,

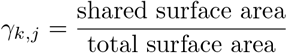

We computed the total surface area as 3.12, and let *A*_*k,j*_ = *A*_*j,k*_ denote the shared (mutually obstructed) surface area of *p*_*k*_ and *p*_*j*_. Then,

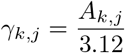

For example, consider two cells who share 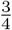 of one of their long sides, as shown in Figure 15a, where the overlap is clearly apparent from the top-side view (see Figure 15b).

Using our model above, we compute the degree of obstruction as 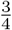 of the area of the long side, divided by the total surface area, like so

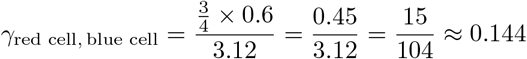

We see, in this example, that the cells obstruct about 14.4% of each other’s surface area. This serves as a simple example model that can be refined later on, such as to account for more realistic cell shapes (especially when the membranes of two cells adhere to each other and thus lead to deviations from the typical ellipsoid-like shape).

## REFERENCES

[1] United Nations. World Population Prospects 2024; 2024. https://population.un.org/wpp/.

[2] World Health Organization. Global cancer burden growing, amidst mounting need for services; 2024. https://www.who.int/news/item/01-02-2024-global-cancer-burden-growing--amidst-mounting-need-for-services.

[3] Hanahan D, Weinberg RA. The Hallmarks of Cancer. Cell. 2000;100(1):57–70. doi:10.1016/S0092-8674(00)81683-9.

[4] Weinberg R. The biology of cancer. Garland Science, Taylor & Francis Group; 2014.

[5] Hanahan D. Hallmarks of Cancer: New Dimensions. Cancer Discovery. 2022;12(1):31–46. doi:10.1158/2159-8290.CD-21-1059.

[6] Alberts B, Bray D, Johnson A, Lewis J, Raff M, Roberts K, et al. Essential Cell Biology. W.W. Norton & Company; 2013.

[7] Folkman J. Tumor angiogenesis: therapeutic implications. New England Journal of Medicine. 1971;285(21):1182–1186.

[8] Kalluri R, Weinberg R. The basics of epithelial-mesenchymal transition. J Clin Invest. 2009;119:1420–1428. doi:10.1172/JCI39104.

[9] Francart M, Lambert J, Vanwynsberghe AM, Thompson EW, Bourcy M, Polette M, et al. Epithelial–mesenchymal plasticity and circulating tumor cells: Travel companions to metastases. Dev Dyn. 2017;247(3):432–450. doi:10.1002/dvdy.24506.

[10] Friedl P, Wolf K. Tumour-cell invasion and migration: diversity and escape mechanisms. Nat Rev Cancer. 2003;5:362–74.

[11] Gupta GP, Massagué J. Cancer Metastasis: Building a Framework. Cell. 2006;127(4):679–695. doi:10.1016/j.cell.2006.11.001.

[12] Lambert AW, Pattabiraman DR, Weinberg RA. Emerging Biological Principles of Metastasis. Cell. 2017;168(4):670–691. doi:10.1016/j.cell.2016.11.037.

[13] Katsaounis D, Harbour N, Williams T, Chaplain MA, Sfakianakis N. A Genuinely Hybrid, Multiscale 3D Cancer Invasion and Metastasis Modelling Framework. Bulletin of Mathematical Biology. 2024;86(6). doi:10.1007/s11538-024-01286-0.

[14] Chaplain MAJ, Lolas G. Mathematical modelling of cancer cell invasion of tissue: the role of the urokinase plasminogen activation system. Math Models Meth Appl Sci. 2005;15:1685–1734.

[15] Chaplain MAJ, Lolas G. Mathematical modelling of cancer invasion of tissue: Dynamic heterogeneity. Net Hetero Med. 2006;1:399–439.

[16] Byrne HM, Chaplain MAJ. Modelling the role of cell-cell adhesion in the growth and development of carcinomas. Mathematical and Computer Modelling. 1996;24(12):1–17.

[17] Burger M, Capasso V, Morale D. On an aggregation model with long and short range interactions. Nonlinear Anal Real World Appl. 2007;8(3):939–958. doi:10.1016/j.nonrwa.2006.04.002.

[18] Morale D, Capasso V, Oelschlager K. An interacting particle system modelling aggregation behavior: from individuals to populations. J Math Biol. 2004;50(1):49–66. doi:10.1007/s00285-004-0279-1.

[19] Scianna M, Preziosi L, Wolf K. A Cellular Potts Model simulating cell migration on and in matrix environments. Mathematical Biosciences and Engineering. 2013;10(1):235–261.

[20] Scianna M, Colombi A. A coherent modeling procedure to describe cell activation in biological systems. CAIM. 2017;8(1):1–22. doi:10.1515/caim-2017-0001.

[21] Van Liedekerke P, Matthias M, Jagiella N, Drasdo D. Simulating tissue mechanics with agent-based models: concepts, perspectives and some novel results. Computational Particle Mechanics. 2015;2(4):401–444.

[22] Van Liedekerke P, Buttenschön A, Drasdo D. Off-lattice agent-based models for cell and tumor growth: numerical methods, implementation, and applications. In: Numerical Methods and Advanced Simulation in Biomechanics and Biological Processes. Springer; 2019.

[23] Anderson A, Chaplain MAJ, Newman E, Steele R, Thompson A. Mathematical Modelling of Tumour Invasion and Metastasis. Comput Math Methods Med. 2000;2:129–154. doi:10.1080/10273660008833042.

[24] Trucu D, Lin P, Chaplain MAJ, Wang Y. A multiscale moving boundary model arising in cancer invasion. Multiscale Modeling & Simulation. 2013;11(1):309–335.

[25] Andasari V, Gerisch A, Lolas G, South AP, Chaplain MAJ. Mathematical modelling of cancer cell invasion of tissue: biological insight from mathematical analysis and computational simulation. J Math Biol. 2011;63(1):141–71.

[26] Sfakianakis N, Chaplain MAJ. Mathematical Modelling of Cancer Invasion: A Review. In: Suzuki T, Poignard C, Chaplain M, Quaranta V, editors. Methods of Mathematical Oncology. vol. 370 of Springer Proceedings in Mathematics & Statistics. Singapore: Springer; 2021. p. 153–172.

[27] Capasso V, Morale D. On the Stochastic Modelling of Interacting Populations. A Multiscale Approach Leading to Hybrid Models. In: Fitzgibbon W, Kuznetsov YA, Neittaanmäki P, Périaux J, Pironneau O, editors. Applied and Numerical Partial Differential Equations: Scientific Computing in Simulation, Optimization and Control in a Multidisciplinary Context. Dordrecht: Springer Netherlands; 2010. p. 59–80.

[28] Colombi A, Scianna M, Preziosi L. Coherent modelling switch between pointwise and distributed representations of cell aggregates. J Math Biol. 2017;74:783–808. doi:10.1007/s00285-016-1042-0.

[29] Carrillo JA, Choi YP, Perez SP. In: A Review on Attractive–Repulsive Hydrodynamics for Consensus in Collective Behavior. Springer International Publishing; 2017. p. 259–298.

[30] Van Liedekerke P, Neitsch J, Johann T, Warmt E, Gonzàlez-Valverde I, Höhme S, et al. A quantitative high-resolution computational mechanics cell model for growing and regenerating tissues. Biomechanics and Modeling in Mechanobiology. 2020;19(1):189–220.

[31] Katsaounis D, Chaplain MAJ, Sfakianakis N. Stochastic differential equation modelling of cancer cell migration and tissue invasion. J Math Biol. 2023;87(1). doi:10.1007/s00285-023-01934-4.

[32] Diez A, Feydy J. Multicellular simulations with shape and volume constraints using optimal transport. arXiv. 2024;.

[33] Chauvière A, Manifacier I, Verdier C, Chagnon G, Cheddadi I, Glade N, et al. A biomechanical model for cell sensing and migration. Computer Methods in Biomechanics and Biomedical Engineering. 2024; p. 1–19.

[34] Chaplain MAJ, Powathil GG. Multiscale Modelling of Cancer Progression and Treatment Control: The Role of Intracellular Heterogeneities in Chemotherapy Treatment. Biophysical Reviews and Letters. 2015;10(02):97–114.

[35] Ramis-Conde I, Drasdo D, Anderson ARA, Chaplain MAJ. Modeling the influence of the E-cadherin–β-catenin pathway in cancer cell invasion: A multiscale approach. Biophysical Journal. 2008;95(1):155–165.

[36] Ramis-Conde I, Chaplain MAJ, Anderson ARA, Drasdo D. Multi-scale modelling of cancer cell intravasation: the role of cadherins in metastasis. Physical Biology. 2009;6(1):016008. doi:10.1088/1478-3975/6/1/016008.

[37] Menci M, Papi M, Porzio MM, Smarrazzo F. On a coupled hybrid system of nonlinear differential equations with a nonlocal concentration. Journal of Differential Equations. 2023;361:288–338. doi:10.1016/j.jde.2023.02.044.

[38] Oliver S, Williams M, Jolly MK, Gonzalez D, Powathil G. Exploring the role of EMT in ovarian cancer progression using a multiscale mathematical model. npj Systems Biology and Applications. 2025;11(1).

[39] Macnamara CK, Caiazzo A, Ramis-Conde I, Chaplain MAJ. Computational modelling and simulation of cancer growth and migration within a 3D heterogeneous tissue: The effects of fibre and vascular structure. Journal of Computational Science. 2020;40:101067.

[40] Chiari G, Delitala ME, Morselli D, Scianna M. A hybrid modeling environment to describe aggregates of cells heterogeneous for genotype and behavior with possible phenotypic transitions. Int J Non-Linear Mech. 2022;144:104063. doi:10.1016/j.ijnonlinmec.2022.104063.

[41] Menci M, Natalini R, Paul T. Microscopic, kinetic and hydrodynamic hybrid models of collective motions with chemotaxis: a numerical study. Mathematics and Mechanics of Complex Systems. 2024;12(1):47–83. doi:10.2140/memocs.2024.12.47.

[42] Jagiella N, Müller B, Müller M, Vignon-Clementel IE, Drasdo D. Inferring growth control mechanisms in growing multi-cellular spheroids of NSCLC cells from spatial-temporal image data. PLoS Computational Biology. 2016;12(2):e1004412.

[43] Sfakianakis N, Madzvamuse A, Chaplain MAJ. A Hybrid Multiscale Model for Cancer Invasion of the Extracellular Matrix. Multiscale Modeling and Simulation. 2020;18(2). doi:10.1137/18M1189026.

[44] Franssen CL, Sfakianakis N, Chaplain MAJ. A novel 3D atomistic-continuum cancer invasion model: In silico simulations of an in vitro organotypic invasion assay. Journal of Theoretical Biology. 2021;522. doi:10.1016/j.jtbi.2021.110677.

[45] Trepat X, Wasserman MR, Angelini TE, Millet E, Weitz DA, Butler JP, et al. Physical forces during collective cell migration. Nature Physics. 2009;5(6):426–430.

[46] Friedl P, Locker J, Sahai E, Segall JE. Classifying collective cancer cell invasion. Nature Cell Biology. 2012;14(8):777–783. doi:10.1038/ncb2548.

[47] Khalil AA, Ilina O, Vasaturo A, Venhuizen JH, Vullings M, Venhuizen V, et al. Collective invasion induced by an autocrine purinergic loop through connexin-43 hemichannels. Journal of Cell Biology. 2020;219(10). doi:10.1083/jcb.201911120.

[48] Janiszewska M, Candido Primi M, Izard T. Cell Adhesion in Cancer: Beyond the Migration of Single Cells. Journal of Biological Chemistry. 2020;295:2495–2505. doi:10.1074/jbc.REV119.007759.

[49] Gerisch A, Chaplain MAJ. Mathematical modelling of cancer cell invasion of tissue: Local and non-local models and the effect of adhesion. J Theor Biol. 2008;250:684–704.

[50] Carrillo JA, Colombi A, Scianna M. Adhesion and volume constraints via nonlocal interactions determine cell organisation and migration profiles. J Theor Biol. 2018;445:75–91.

[51] Colombi A, Falletta S, Scianna M, Scuderi L. An integro-differential non-local model for cell migration and its efficient numerical solution. Math Comput Simul. 2021;180:179–204.

[52] Painter KJ, Bloomfield JM, Sherratt JA, Gerisch A. A Nonlocal Model for Contact Attraction and Repulsion in Heterogeneous Cell Populations. Bull Math Biol. 2015;77(6):1132–1165. doi:10.1007/s11538-015-0080-x.

[53] Conte M, Loy N. A Non-Local Kinetic Model for Cell Migration: A Study of the Interplay Between Contact Guidance and Steric Hindrance. SIAM J Appl Math. 2023;84(3):S429–S451. doi:10.1137/22m1506389.

[54] Falcó C, Baker RE, Carrillo JA. A Local Continuum Model of Cell-Cell Adhesion. SIAM Journal on Applied Mathematics. 2024;84(3):S17–S42. doi:10.1137/22M1506079.

[55] D’Orsogna MR, Chuang YL, Bertozzi AL, Chayes LS. Self-Propelled Particles with Soft-Core Interactions: Patterns, Stability, and Collapse. Phys Rev Lett. 2006;96(10). doi:10.1103/physrevlett.96.104302.

[56] Belyaev AV, Fedotova IV. Molecular mechanisms of catch bonds and their implications for platelet hemostasis. Biophysical Reviews. 2023;15(5):1233–1256. doi:10.1007/s12551-023-01144-8.

[57] Zhigun A, Rajendran ML. Modelling non-local cell-cell adhesion: a multiscale approach. Journal of Mathematical Biology. 2024;88(5). doi:10.1007/s00285-024-02079-8.

[58] Viji Babu PK, Mirastschijski U, Belge G, Radmacher M. Homophilic and heterophilic cadherin bond rupture forces in homo- or hetero-cellular systems measured by AFM-based single-cell force spectroscopy. European Biophysics Journal. 2021;50(3–4):543–559. doi:10.1007/s00249-021-01536-2.

[59] Loh CY, Chai J, Tang T, Wong W, Sethi G, Shanmugam M, et al. The E-Cadherin and N-Cadherin Switch in Epithelial-to-Mesenchymal Transition: Signaling, Therapeutic Implications, and Challenges. Cells. 2019;8(10):1118. doi:10.3390/cells8101118.

[60] Rakshit S, Zhang Y, Manibog K, Shafraz O, Sivasankar S. Ideal, catch, and slip bonds in cadherin adhesion. Proceedings of the National Academy of Sciences. 2012;109(46):18815–18820. doi:10.1073/pnas.1208349109.

[61] Sivasankar S. Tuning the Kinetics of Cadherin Adhesion. Journal of Investigative Dermatology. 2013;133(10):2318–2323. doi:10.1038/jid.2013.229.

[62] Panorchan P, Thompson MS, Davis KJ, Tseng Y, Konstantopoulos K, Wirtz D. Single-molecule analysis of cadherin-mediated cell-cell adhesion. Journal of Cell Science. 2006;119(1):66–74. doi:10.1242/jcs.02719.

[63] Dash S, Duraivelan K, Hansda A, Kumari P, Chatterjee S, Mukherjee G, et al. Heterophilic recognition between E-cadherin and N-cadherin relies on same canonical binding interface as required for E-cadherin homodimerization. Archives of Biochemistry and Biophysics. 2022;727:109329. doi:10.1016/j.abb.2022.109329.

[64] MATLAB. MATLAB Version 25.1.0.2943329 (R2025a). Natick, Massachusetts: The Mathworks, Inc.; 2025.

[65] Bezanson J, Edelman A, Karpinski S, Shah VB. Julia: A fresh approach to numerical computing. SIAM Review. 2017;59(1):65–98.

[66] Shashni B, Ariyasu S, Takeda R, Suzuki T, Shiina S, Akimoto K, et al. Size-Based Differentiation of Cancer and Normal Cells by a Particle Size Analyzer Assisted by a Cell-Recognition PC Software. Biological and Pharmaceutical Bulletin. 2018;41(4):487–503. doi:10.1248/bpb.b17-00776.

